# Regulation of lung cancer initiation and progression by the stem cell determinant Musashi

**DOI:** 10.1101/2024.04.11.589053

**Authors:** Alison G. Barber, Cynthia M. Quintero, Michael Hamilton, Nirakar Rajbhandari, Roman Sasik, Yan Zhang, Carla F. Kim, Hatim Husain, Xin Sun, Tannishtha Reya

## Abstract

Despite advances in therapeutic approaches, lung cancer remains the leading cause of cancer-related deaths. To understand the molecular programs underlying lung cancer initiation and maintenance, we focused on stem cell programs that are normally extinguished with differentiation but can be reactivated during oncogenesis. Here we have used extensive genetic modeling and patient derived xenografts to identify a dual role for Msi2: as a signal that acts initially to sensitize cells to transformation, and subsequently to drive tumor propagation. Using Msi reporter mice, we found that Msi2-expressing cells were marked by a pro-oncogenic landscape and a preferential ability to respond to Ras and p53 mutations. Consistent with this, genetic deletion of Msi2 in an autochthonous Ras/p53 driven lung cancer model resulted in a marked reduction of tumor burden, delayed progression, and a doubling of median survival. Additionally, this dependency was conserved in human disease as inhibition of Msi2 impaired tumor growth in patient-derived xenografts. Mechanistically, Msi2 triggered a broad range of pathways critical for tumor growth, including several novel effectors of lung adenocarcinoma. Collectively, these findings reveal a critical role for Msi2 in aggressive lung adenocarcinoma, lend new insight into the biology of this disease, and identify potential new therapeutic targets.

## Introduction

Lung cancer is the second most common type of cancer. While the death rate due to lung cancer has decreased over time due to a drop in the prevalence of smoking, it is still by far the leading cause of cancer-related deaths in men and women, resulting in more deaths than colon, breast, and prostate cancer combined^1^. Non-small-cell lung cancer (NSCLC) is the most prevalent form of lung cancer, accounting for 84% of lung cancer cases. Within this cancer type, adenocarcinoma is the most frequent subtype, accounting for 50% of all NSCLCs^2,3^. Most cases of lung cancer are diagnosed at late-stage— when there is regional or distant spread of the disease—at which time treatment options are limited^4^. Despite advances in lung cancer therapy, the 5-year survival rates for late-stage (i.e. regional or distant) NSCLC are 37% to 9%, respectively^5^. These survival rates highlight the critical need for improved understanding of the biology of this disease in order to develop more effective treatments.

To understand the key dependencies of lung cancer, we have focused on stem cell programs which are often co-opted to perpetuate an aggressive, undifferentiated state. The stem cell fate determinant Musashi-2 (Msi2) is one such signal that has been shown to be critical in development as well as advanced cancers^6,7,8^; however, its role in lung cancer is not well understood. Msi2 has been reported to be overexpressed in human lung adenocarcinoma and regional metastases, and its inhibition in human cell lines has been shown to reduce invasive and metastatic potential *in vitro*^9,10^. However, this study focused on *in vitro* analysis utilizing human NSCLC cell lines; thus, no evidence for a role of Msi2 *in vivo* using definitive genetic mouse models exist. Further, whether Msi2 is a dependency for tumor initiation or required for continued propagation, and which pathways are regulated by Msi2 to drive tumor growth, also remain unknown. Here we have utilized a combination of genetic models and patient-derived xenografts to determine the role of Msi2 signaling in the growth and progression of lung adenocarcinoma. Using Msi2 knock-in reporter mice and Msi2 conditional knockout mice, we show that Msi2 plays a dual role in lung adenocarcinoma, acting as a signal that not only is essential for initiation, but one that continues to be required post-establishment in both genetically-engineered mouse models and in patient-derived xenografts. Finally, using RNA-sequencing analysis we show that Msi2 can trigger a range of pathways critical for tumor growth, including several new effectors of lung adenocarcinoma. These findings suggest that Msi2 is a critical regulator of lung adenocarcinoma and offer new insight into the signals that facilitate transformation and support disease progression.

## Results

### Msi2 is expressed in normal lung and in adenocarcinoma

To determine whether Musashi regulates the growth and progression of aggressive lung cancer we first determined whether Msi2 is expressed in stem cells of the distal lung. We used a Msi2^eGFP/+^ knock-in reporter mouse to track endogenous Msi2 expression^6^, and found that Msi2 is expressed in 39% of Lin^-^EpCAM^+^ lung epithelial cells (Fig. 1A) and, more specifically, in 37% of cells enriched for Club/BASC cells and 26% of cells enriched for AT2 cells—the known stem cell populations of the distal lung (Fig. 1B-C). Interestingly, although Msi2 expression could only be detected in a fraction of distal lung stem cells, widespread expression of Msi2 was often observed in lung adenocarcinomas formed in Kras^G12D/+^; p53^fl/fl^ mice (Fig. 1D). The expansion of Msi2 expression in lung tumors compared to normal lung epithelia suggests that normal cells expressing Msi2 are more sensitive to transformation, or that once transformed, Msi2 expression is amplified in tumor cells.

**Figure 1:**
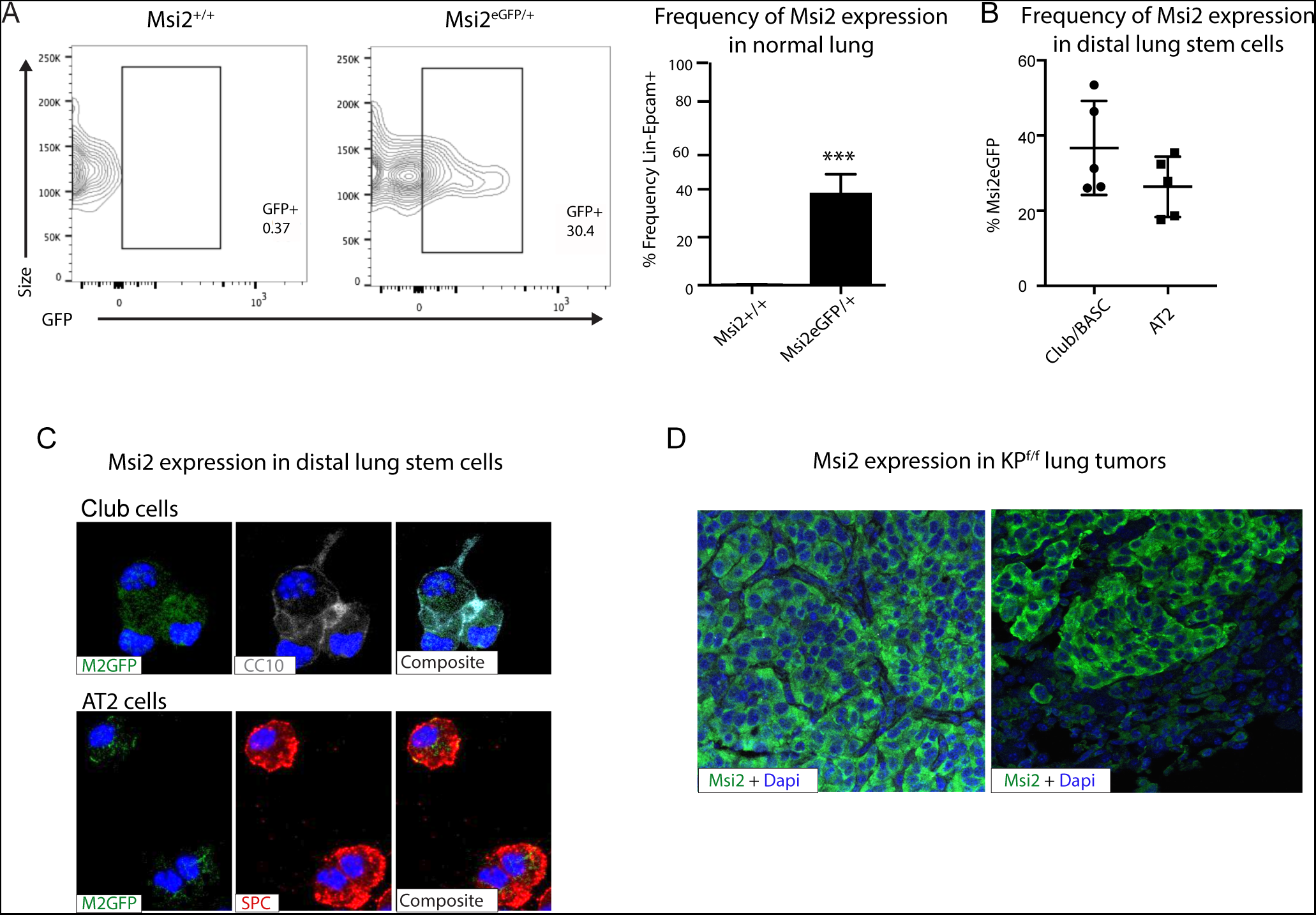
Msi2 is expressed in the stem cells of the distal lung and in lung adenocarcinoma. A) Msi2 is expressed in the Lin^-^EpCAM^+^ lung epithelial cells of Msi2^GFP/+^ (M2GFP) reporter mice. Representative FACS plots shown for non-reporter (Msi2^+/+^, left) and M2GFP reporter mouse lungs (Msi2^eGFP/+^middle). Frequency of M2GFP-expressing cells in lung epithelia (right; n = 2 non-reporter, n = 5 reporter). Data represented as mean ± SD, one outlier identified and removed using the Grubb’s test, ***p < 0.001 by Student’s t-test. B-C) Msi2 is expressed in the known stem cell populations of the distal lung. B) Frequency of Msi2 expression in the Lin^-^EpCAM^+^Sca1^+^ enriched Club/BASC and the Lin^-^EpCAM^+^Sca1^-^ enriched AT2 cell populations (n=5). C) Representative images of cytospins from M2GFP reporter mouse Lin^-^EpCAM^+^ lung epithelial cells showing expression of Msi2 expression in Club cells (top row, marked by CC10 staining) and AT2 cells (bottom row, marked by SPC staining). D) Representative images of Msi2 in tumors from KP^f/f^ mice; some tumors display ubiquitous expression of Msi2 (left), while others display more heterogeneous expression (right).

### Msi2-expressing cells are more responsive to transformation by mutant Ras and p53

To determine whether Msi2 expression marks cells with increased capacity for tumor growth, we generated Msi2-reporter Kras^G12D/+^; p53^fl/fl^ mice with inducible Cre expression (Msi2^eGFP/+^; Kras^G12D/+^; p53^fl/fl^; RosaCre^ER/+^; Fig. 2A), delivered tamoxifen, isolated Msi2^+^ and Msi2^-^ lung epithelial cells, and tested sphere-forming capacity. As shown in Fig. 2B, Msi2^+^ cells had enhanced sphere-forming capacity, with a 6-fold increase in sphere-forming observed at primary passage, and this capacity was retained for up to 4 passages (Fig. 2B). Further, spheres that were isolated and transplanted subcutaneously into the flanks of immunocompromised mice developed Msi2-expressing tumors *in vivo* as well as metastatic lesions in the lung, demonstrating that the observed sphere-forming capacity is reflective of tumorigenicity and not merely stem cell capacity (Fig. 2C-E). To understand the basis for the enhanced tumorigenic capacity of Msi2^+^ cells, we carried out RNA-sequencing (RNAseq) to examine the transcriptional landscape of Msi2-expressing cells in normal lung epithelia. Interestingly, the differentially expressed gene (DEG) profile showed that Msi2^+^ cells are transcriptionally distinct from Msi2^-^ cells (Fig. 2F). Gene Set Enrichment Analysis (GSEA) further revealed that Msi2^+^ cells are significantly enriched for signatures related to developmental and stem cell signaling, consistent with the known role of Msi2 as a critical regulator of the stem cell state^11,12,13^, as well as signatures related to oncogenic signaling and therapy resistance (Fig. 2G, 2I, 2K) which are associated with aggressive cancers. Over 1,900 differentially expressed genes were found to be enriched in the Msi2^+^ population. In line with the GSEA results, these enriched genes included those involved in developmental and stem cell signaling (such as *Dll1, Jag2,* and *Notch3)* ^14–17^, oncogenic signaling (such as *Akt3, Ret, Myb, and Fos*)^18–21^, and therapy resistance (such as *Axl*, *Egr1,* and several glutathione S transferases implicated in platinum drug resistance including *Gstp1 and Gsta2*)^22,23,24^ (Fig. 2H, 2J, 2L). These findings suggest that prior to tumor initiation, the transcriptional landscape of Msi2-expressing cells is enriched in signaling programs that are conducive to tumorigenesis, which may make them uniquely poised for transformation.

**Figure 2.**
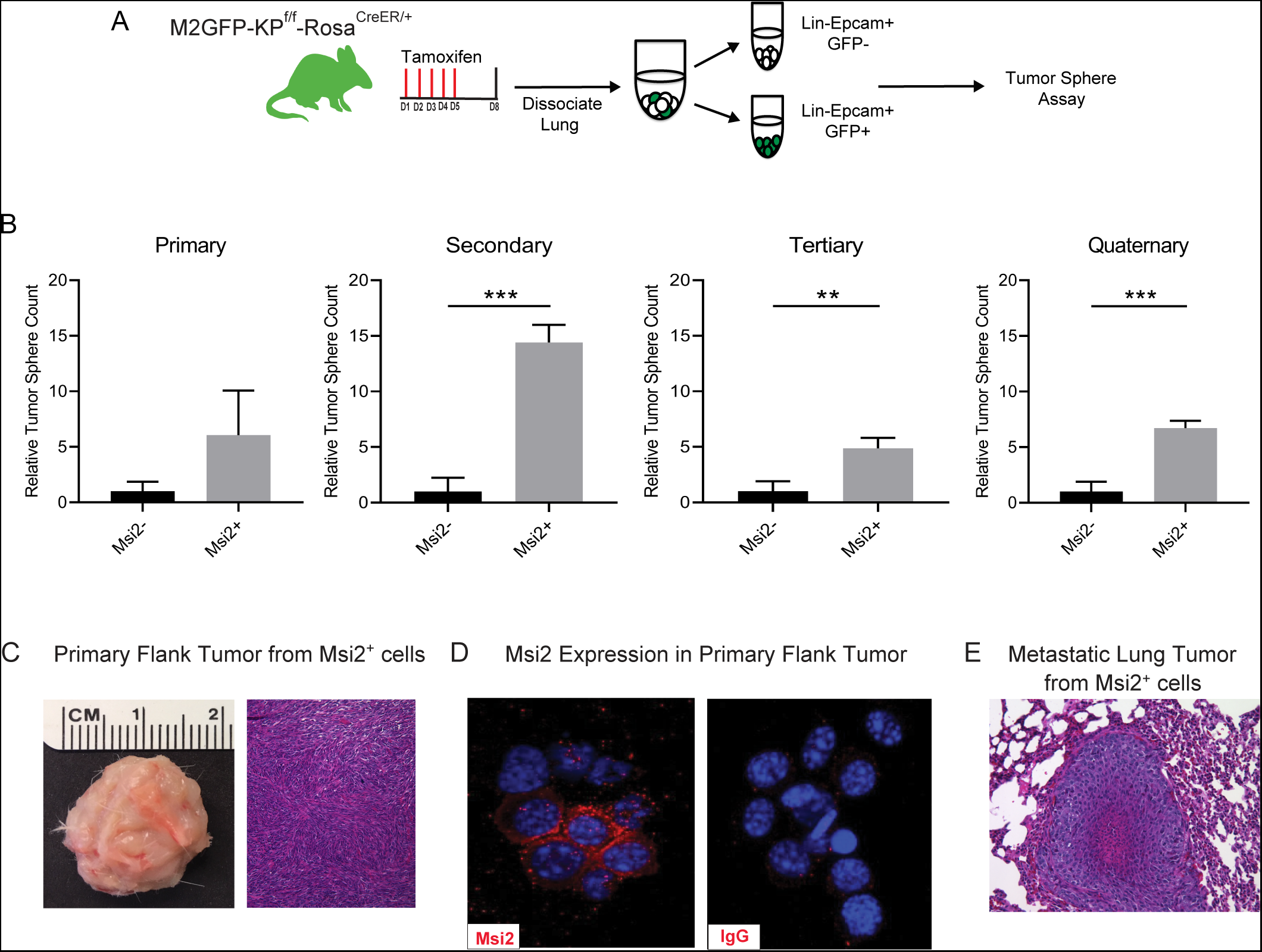
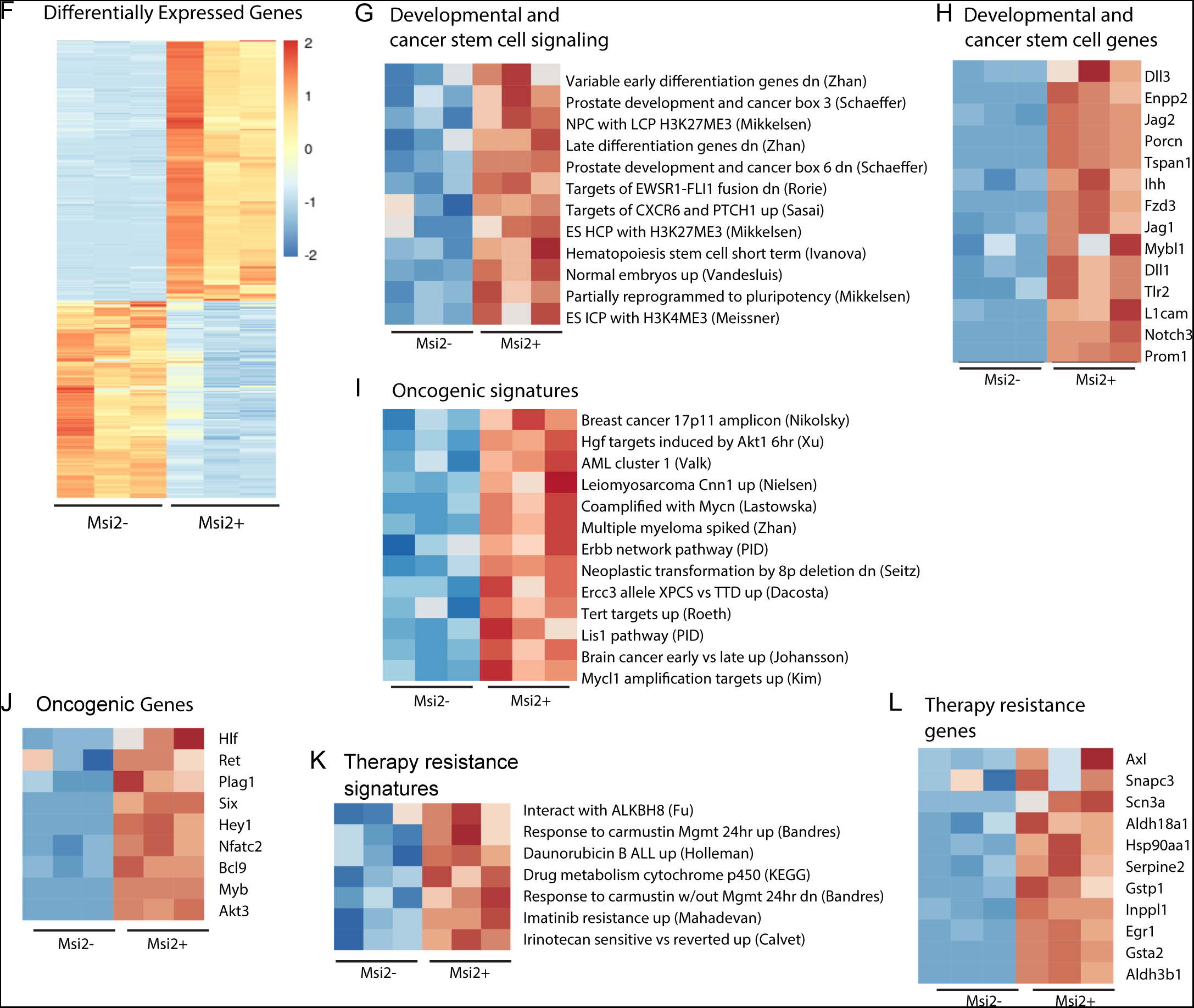
Msi2-expressing cells preferentially favor tumor growth in lung adenocarcinoma. A) Schematic of the strategy used to measure the tumor sphere forming capacity of Msi2-expressing and non-expressing cells *in vitro*. Msi2GFP/+; KrasG12D/+; p53fl/fl; RosaCreER/+ (M2GFP-KPf/f -RosaCreER/+) mice were treated with tamoxifen for 5 days to induce recombination of floxed alleles. Three days following the final dose of tamoxifen, lungs were dissociated and Msi2-expressing (GFP+) or non-expressing (GFP-) cells were isolated by FACS and then plated in a tumor sphere assay. B) Msi2-expressing cells isolated from KPf/f lung epithelia following tamoxifen treatment preferentially form tumor spheres over multiple cell passages *in vitro* as compared to non-expressing cells. Representative experiment shown, (n = 3). Data represented as mean ± SD, **p<0.01, ***p<0.001 by Student’s t-test. C-E) Tumor spheres formed by Msi2+ cells *in vitro* form aggressive tumors *in vivo*. Tumor spheres isolated from Msi2+ cells after quaternary passage *in vitro* and transplanted *in vivo* form highly aggressive flank tumors (C) that retain Msi2 expression (D) and are able to metastasize to the lung (E). F) Heatmap showing significant differentially expressed genes between Msi2-expressing cells and non-expressing cells. G-L) Gene Set Enrichment Analysis of Msi2-expressing normal lung epithelial cell gene signatures. Heatmaps of gene signatures and selected genes that are involved in developmental and stem cell signaling (G-H), oncogenic signaling (I-J), and therapy resistance (K-L). Red represents gene signatures or genes that are upregulated in the presence of Msi2, while blue represents gene signatures or genes that are downregulated in the absence of Msi2.

### Genetic deletion of Msi2 leads to a decrease in tumor incidence, burden, and progression

To determine if Msi2 not only marks a cell with an enhanced capacity for lung adenocarcinoma growth, but whether it may also be required for initiation of these tumors, we compared wild type and Msi2 knock-out Kras^G12D/+^; p53^fl/fl^ mice (Msi2^-/-^; Kras^G12D/+^; p53^fl/fl^) (Fig. 3A). In this model, the *Msi2* gene has been disrupted via gene trap mutagenesis^7^, while the inhalation of adenoviral-Cre (Ad-Cre) results in the conditional activation of Kras and inactivation of p53 in the lung^25^. Importantly, loss of Msi2 led to the formation of significantly fewer (84%) tumor lesions as well as a significantly lower (85%) overall tumor burden in the lung at a 14-week midpoint (Fig. 3B-D). Further, while there was no difference in the frequency of low-grade tumors, there was a significant decrease in the frequency of mid- and high-grade tumors (45-54%), suggesting that loss of Msi2 delayed tumor progression from low to high grade (Fig. 3E). Importantly, Msi2^-/-^ KP^f/f^ mice infected with Ad-Cre developed tumors less frequently than their wild type counterparts (61% Msi2^-/-^ vs 83% Msi2^+/+^; Fig. 3F). To confirm that the population of lung stem cells is not reduced in Msi2 knockout mice and therefore contributing to a decrease in tumor lesions, we stained the lungs of WT and Msi2 knockout mice for Club/BASC and AT2 lung stem cells and saw no difference (data not shown). Additionally, Msi2^-/-^ KP^f/f^ mice displayed a significant increase in overall survival compared to Msi2^+/+^ KP^f/f^ mice. Msi2^+/+^ KP^f/f^ mice had a median survival of 209 days while the Msi2^-/-^ KP^f/f^ mice had a median survival of 428 days, reflecting a 2-fold increase in survival (p = 0.04, Fig. 3G). Notably, the majority of lung tumors that formed in Msi2^-/-^ KP^f/f^ mice were found to express Msi2, suggesting that the tumors that formed in Msi2^-/-^ KP^f/f^ mice were escapers that re-expressed Msi2, underscoring the dependency on Msi2 signaling for tumor formation (Fig. 3H). Taken together these results suggest that Msi2 not only marks cells with an enhanced capacity for tumor growth but is also required for initiation and progression of lung adenocarcinoma.

**Figure 3:**
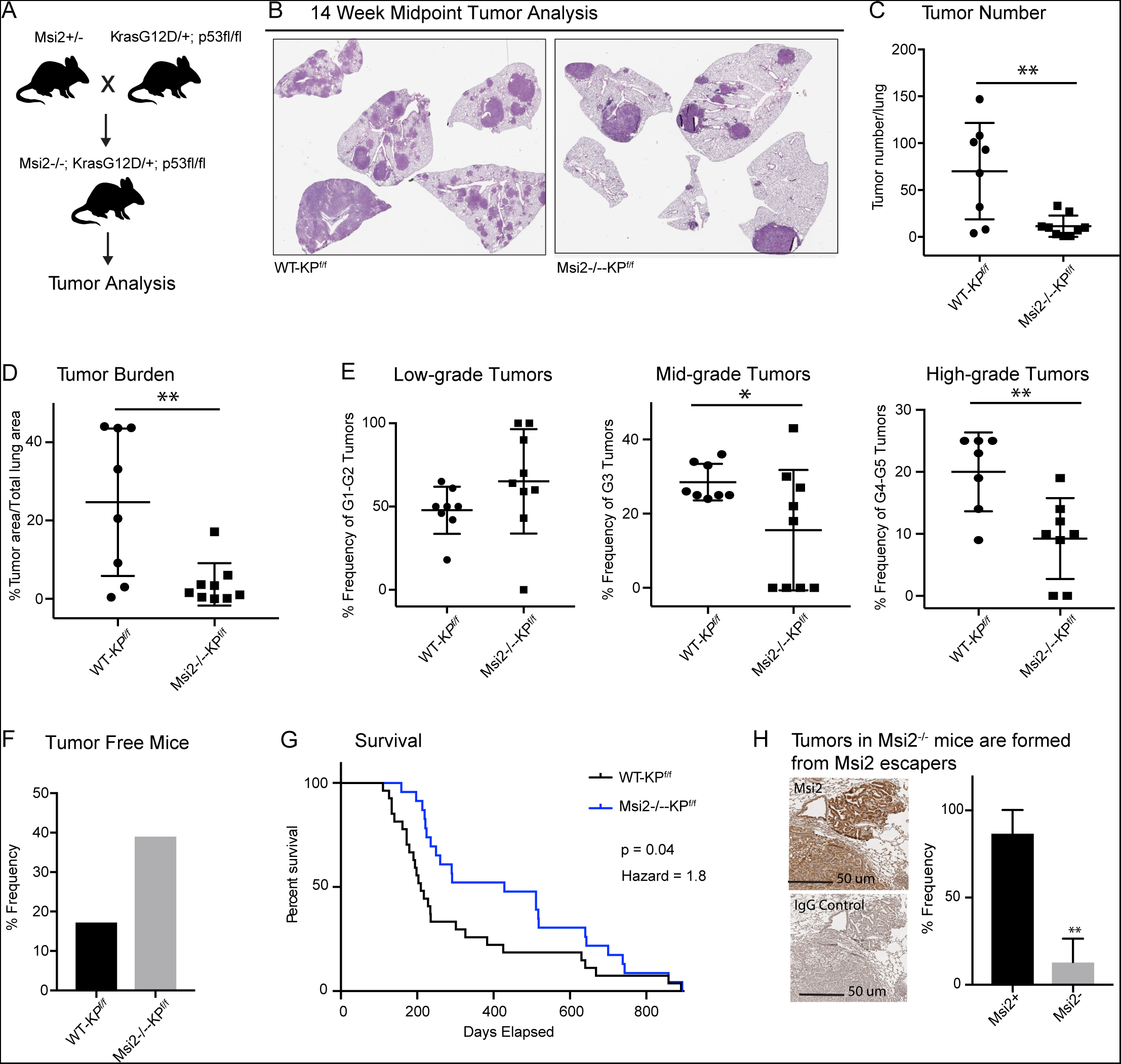
Msi2 is required for lung adenocarcinoma growth and progression. A) Breeding scheme used to generate the Msi2^-/-^; Kras^G12D/+^; p53^fl/fl^ (Msi2^-/-^-KP^f/f^) model of lung adenocarcinoma. B) Representative images of wild type (left) and Msi2^-/-^ (right) Kras^G12D/+^; p53^fl/fl^ lungs gender and age matched mice at 14 weeks after tumor initiation. C-D) The number of tumors formed (C) and overall tumor burden (D) is significantly reduced in Msi2^-/-^ KP^f/f^ mice. E) Msi2^-/-^ KP^f/f^ mice have a significant reduction in the frequency of mid- and high-grade tumors. Data represented as mean ± SD, two outliers identified and removed for high-grade tumors using the Grubb’s test *p < 0.05, **p<0.01 by Student’s t-test. F) The tumor take rate is reduced in Msi2^-/-^ KP^f/f^ mice. While only 17% of AdCre-infected wild ype mice are tumor-free at the time of death, more than twice as many (39%) Msi2^-/-^ KP^f/f^ mice are tumor-ree at the time of death. G) Msi2^-/-^ KP^f/f^ have significantly increased survival (p = 0.04, hazard ratio = 1.8; n = 27 WT-KP^f/f^, n = 23 Msi2^-/-^-KP^f/f^), with a median survival of 428 days compared to 209 days for wild type mice. Log-rank test was used to determine the difference in survival curves between wild type and Msi2^-/-^ KP^f/f^ mice. H) Lung tumors in Msi2^-/-^ KP^f/f^ mice express Msi2. Representative immunohistochemical staining (left) for Msi2 (top) or IgG control (bottom) in lung tumors from a 16-week old Msi2^-/-^ KP^f/f^ mouse. Quantification of the frequency of Msi2-expressing tumors from Msi2^-/-^ KP^f/f^ mice shows hat while some tumors lack Msi2 expression the majority of tumors express Msi2 (right, n = 3). Data represented as mean ± SD, **p<0.01 by Student’s t-test.

The analyses described above were performed in Msi2^-/-^ KP^f/f^ mice, and therefore in an environment in which Msi2 is absent at the onset of tumorigenesis. To test whether established cancers have a similar dependency on Msi2 for their growth, we knocked down Msi2 in primary tumor cell lines generated from Kras^G12D/+^; p53^fl/fl^ lung tumors (KP^f/f^ cell line) and measured tumorsphere formation. Importantly, loss of Msi2 led to a significant decrease (75%) in tumorsphere formation *in vitro*, suggesting that Msi2 is required for tumor cell propagation (Fig. 4A-B). To confirm this finding *in vivo*, Msi2 knockdown KP^f/f^ cells were transplanted into the flanks of syngeneic mice, and tumor growth monitored over time. As shown in Figure 4, the Msi2-knockdown tumors had significantly reduced (89%) tumor cell numbers (Fig. 4C-D), indicating that established tumors remain dependent on Msi2 signaling for growth. To determine the relevance of our findings for human disease, we examined the effect of Msi2 inhibition on primary patient samples. To this end, patient samples were transplanted into immunocompromised mice to generate patient-derived xenografts (PDXs). Once established, PDX tumors were harvested, dissociated, transduced with either shControl or shMsi2 lentivirus, and subsequently transplanted into the flanks of immunocompromised mice (Fig. 4E). Importantly, although an equivalent number of shControl and shMsi2 cells were transplanted into recipient mice, there was a notable delay in the growth of Msi2-knockdown tumors, which were 64-88% smaller than control tumors at endpoint (Fig. 4F, Supplementary Fig. 1). These findings suggest that human lung adenocarcinomas are dependent on Msi2 for their continued growth.

**Figure 4:**
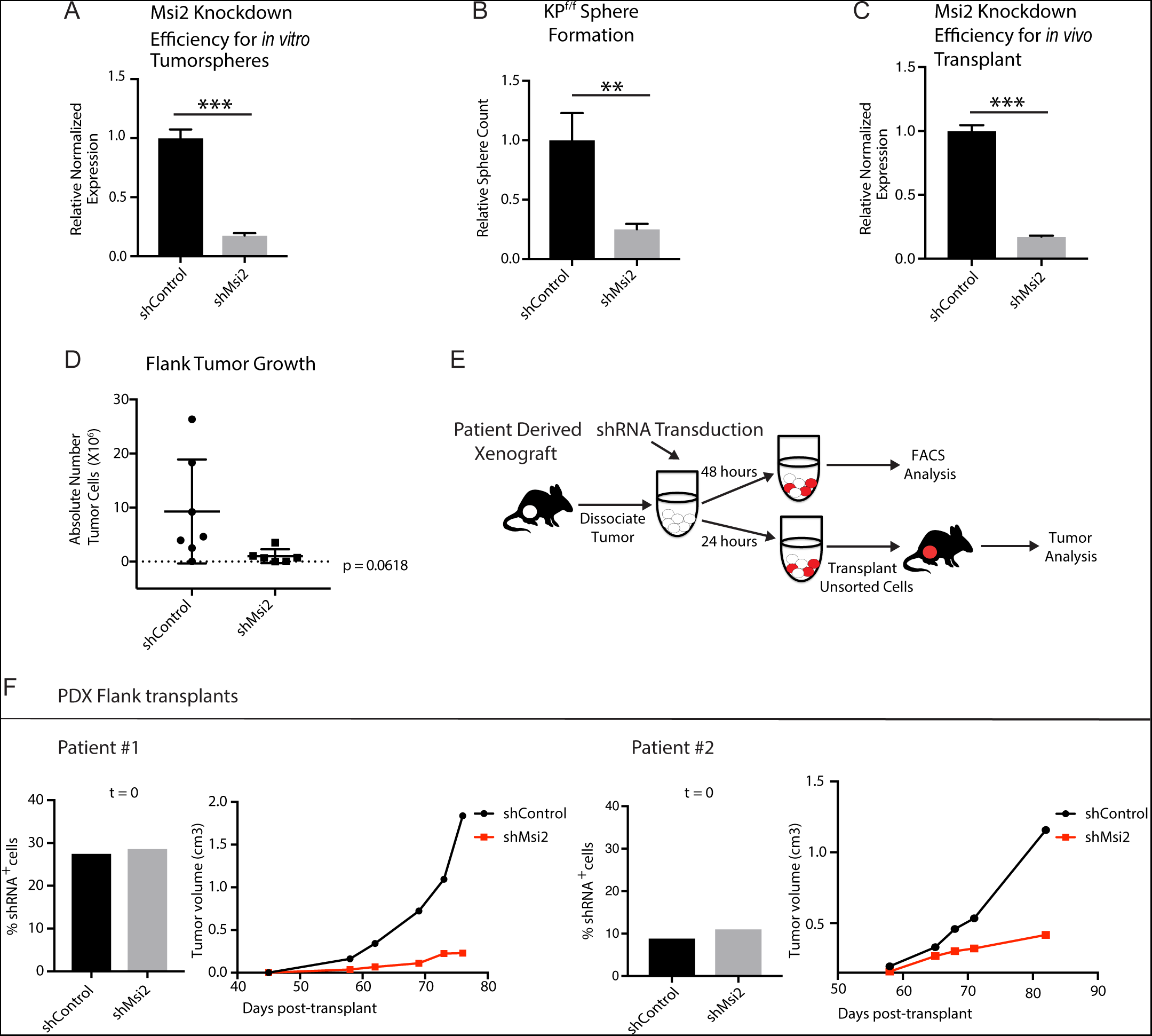
Loss of Msi2 impairs growth of established cancer. A) Mouse KP^f/f^ cells transduced with shMsi2 and used for *in vitro* tumorsphere assays have an 83% reduction in *Msi2* mRNA expression. B) Loss of *Msi2* significantly impairs tumor sphere formation *in vitro* (n =3). C) Mouse KP^f/f^ cells transduced with shMsi2 and used for *in vivo* flank transplant assays have an 83% reduction in *Msi2* mRNA expression. D) Loss of *Msi2* consistently impairs tumor growth in flank transplants *in vivo*. Data represented as mean ± SD, one outlier identified and removed using the Grubb’s test **p<0.01 by Student’s t-test. E) Schematic for testing the impact of Msi2 loss on growth of patient-derived xenografts *in vivo*. Patient-derived xenografts were harvested, dissociated, transduced with shMsi2 or shControl virus and split into two pools that were incubated for either 24 or 48 hours. After 24 hours, one pool of cells (containing a mixture of transduced and untransduced cells) was transplanted into the flanks of NSG mice. After 48 hours the remaining pool of cells was analyzed via FACS to determine the frequency of cells infected with shRNA. F) The frequency of infection was comparable for shControl and shMsi2 in two independent patient samples. Reduced *Msi2* expression led to pronounced inhibition of tumor growth in two independent patient-derived xenografts.

### Msi2 regulates signaling pathways critical for tumor growth

To understand the mechanism by which Msi2 impacts tumor growth, we knocked down Msi2 in KP^f/f^ cell lines and carried out an RNAseq analysis comparing control and knock-down cells. Principal component analysis shows a clear separation between the control and Msi2-knockdown cells and significant changes in over ∼170 genes, highlighting the marked impact that loss of Msi2 has on the transcriptional profile of lung adenocarcinoma (Fig. 5A). Consistent with the role of Msi2 in stem cell maintenance, the inhibition of Msi2 in lung cancer cells had a significant impact on developmental and stem cell signaling programs and genes, this included *Porcn,* the transcription regulator *Nupr1,* and the NuRD complex member *Mbd3*, which have been shown to play a role in the maintenance of the stem cell state (Fig. 5B-C)^26–29^. In addition, Msi2 impacted a broad range of other signaling programs and their associated genes, including DNA repair and metabolism. DNA repair genes included *Brca1*, *Atm*, and several Fanconi anemia complementation group genes^30,31,32^, and metabolism programs included the gluconeogenesis enzyme *Pck2*, which has been shown to promote lung cancer cell survival in low glucose conditions, several isoforms of *Cpt1*, a key regulator in fatty acid oxidation known to promote tumor growth in a variety of cancers, and the glutamate dehydrogenase *Glud1*, which has been shown to promote tumorigenesis in lung cancer xenografts (Fig. 5D-G^33,34,35^). Interestingly, several known regulators of lung adenocarcinoma such as *Gli1*, a canonical downstream effector of the Hedgehog signaling pathway, βIII tubulin (*Tubb3)*, and CC Chemokine Receptor 1 (*Ccr1)* were found to be significantly downregulated following Msi2 inhibition as well (Fig. 5H)^36–44^. These data suggest that Msi2 is a regulator of a variety of signaling pathways that are critical to tumor growth and lung cancer progression. In addition to these known regulators, a number of genes not previously implicated in lung cancer were found to be significantly downregulated following Msi2 inhibition, suggesting that they may play a role in tumor growth (Fig. 5I-J). To identify potential novel regulators of lung adenocarcinoma we focused on significantly downregulated genes that were highly expressed, which narrowed our gene list to 30 potential candidates. PCR based validation and prioritization of those genes with no known role in lung cancer led to a subset that included genes such as the E3 ubiquitin ligase *Rnf157*, which promotes neuronal growth through the inhibition of apoptosis^45^, Synaptotagmin 11 (*Syt11)*, a critical mediator of neuronal vesicular trafficking^46^, prostoglandin D2 synthase (*Ptgds*) known to be involved in reproductive organ development^47^, and ADP ribosylation like binding protein (*Arl2bp),* a known regulator of STAT3 signaling^48^. To test whether these genes regulate tumor cell growth we knocked down each of these genes (*Ptgds*, *Arl2bp*, *Rnf157*, and *Syt11*) in the KP^f/f^ cell line. Inhibition of each target led to a significant decrease (88-98%) in the formation of tumor spheres *in vitro* (Fig. 5K). These findings not only identify these genes as downstream effectors of Msi2 activity but more generally show that this dataset could be an important resource to identify new regulators of lung adenocarcinoma growth.

**Figure 5:**
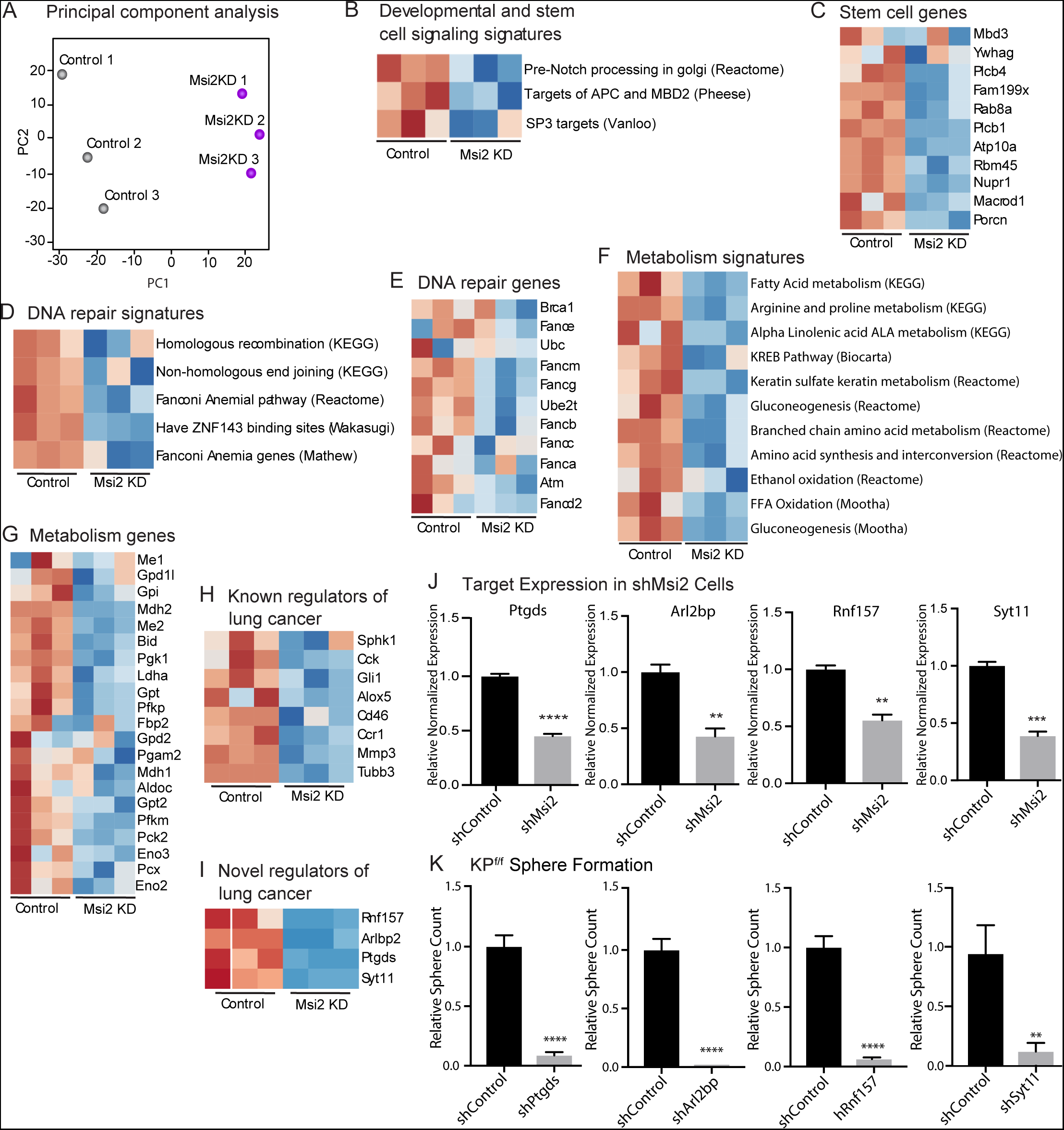
Msi2 regulates oncogenic signaling in lung adenocarcinoma. A) Principal component analysis of KP cells transduced with shMsi2 (Msi2KD, purple) or shControl (gray). BG) Gene Set Enrichment Analysis of Msi2KD gene signatures. Heatmaps of gene signatures and selected genes that are involved in developmental and stem cell signaling (B-C), DNA repair (D-E), and metabolism (F-G) and are downregulated (shown in blue) following the loss of Msi2. H-I) Known (H) and novel (I) regulators of lung cancer that are downregulated following loss of Msi2. B-I) Red represents genes or gene signatures that are upregulated in the presence of Msi2, while blue represents genes that are downregulated following the loss of Msi2. J) Confirmation of downregulated genes in Msi2KD cells using qRT-PCR analysis. K) *In vitro* functional analysis of novel effectors of lung cancer (Ptgds, Arl2bp, Rnf157, Sty11) downregulated by Msi2. KP_f/f_ cells were transduced with shRNA to inhibit the genes of interest and analyzed for the resulting impact on tumor sphere formation *in vitro*. Sphere formation, n=3 per condition. Data represented as mean ± SD,**p<0.01, ***p<0.001, ****p<0.0001 by Student’s t-test.

## Discussion

The stem cell fate determinant Msi2 is commonly found to be expressed in carcinomas^10,49,50,51^ and Msi signaling has been shown to contribute to colon adenomas^52^ and maintain the aggressive, undifferentiated state in hematologic malignancies and pancreatic cancer^7,6^. Here we have used genetically-engineered and autochthonous models to show that Msi2 is critically required in the development and progression of primary lung adenocarcinoma. Msi2 signaling emerges as a key dependency at initiation and continues to be needed during tumor progression. Using the well-established Kras^G12D^; p53^flox/flox^ (KP^fl/fl^) mouse model of lung adenocarcinoma^53^, we created a Msi2 knock-out lung adenocarcinoma model and found that deletion of Msi2 at initiation significantly impaired tumor incidence, reduced tumor burden and progression, and cumulatively led to a doubling of median survival. Furthermore, loss of Msi2 impaired propagation of established KP^fl/fl^ tumors *in vitro* and *in vivo*. In order to determine whether these findings were relevant for the human disease, we used patient-derived xenografts, which have proven to be a highly valuable tool for understanding the biology of human cancers and have been shown to faithfully predict therapeutic response in a majority of patients^54,55,56^. Inhibition of Msi2 in PDXs also led to impaired tumor growth *in vivo*, suggesting that dependence on Msi2 is conserved in human lung adenocarcinoma.

Our finding that the loss of Msi2 prior to transformation significantly impacts tumor initiation prompts the interesting question of whether Msi2 signaling can influence early tumor growth for lung adenocarcinoma. Previous use of the KP^fl/fl^ model has illustrated how both the cellular and molecular context at the time of transformation influences subsequent cancer development^57^. Using cell-type specific Ad-Cre to drive the expression of either Kras^G12D^ alone or Kras^G12D^ in combination with p53^flox/flox^, work by Sutherland et al.^57^ has shown that the cell of origin for adenocarcinoma differs based on the genetic mutations used to drive the cancer. When Kras^G12D^ alone was used as the driver, AT2 cells were able to form adenomas which then progressed to adenocarcinomas, while Club cells were only able to form adenomas which did not progress further. However, when Kras^G12D^ was used in combination with p53^flox/flox^, both cell types were capable of forming adenomas which could then progress to adenocarcinomas. While both genetic drivers and the cell of origin are key components for tumor formation, whether there are signaling factors that endow the cell of origin with the capacity to transform when presented with these key drivers is an area that has been less explored. In this context, our work sheds new light on such signals required at initiation as we found that Msi2 is expressed in a subpopulation of distal lung stem cells, that cells expressing Msi2 are preferentially sensitive to transformation, and that its genetic ablation leads to a marked decrease in tumor initiation and progression. The results of our RNAseq analysis suggest that this enhanced capacity may be established through an enrichment in signaling programs involved in the maintenance of an undifferentiated state as well as those known to support aggressive tumor growth which then provides a permissive environment for early tumor growth and development. These findings identify a new role for Msi2 signaling as an enhancer of tumor growth for lung adenocarcinoma and indicate that subpopulations harboring these signals may be particularly primed for transformation.

The continued dependency on Msi2 signaling following tumor initiation illustrates the potency of Msi2 as a regulator of tumor progression; thus, defining the mechanism by which Msi2 exerts this influence in lung adenocarcinoma would be critical to understanding the molecular basis of its broad influence. Here, our RNAseq analysis suggests that Msi2 may act in part as a regulator of the stem cell state. Interestingly, this analysis also showed that the role of Msi2 may extend to the regulation of processes crucial for tumor growth, such as metabolism and DNA repair, thus placing it upstream of multiple potent oncogenic signals. This work also led to the identification of several novel effectors of lung adenocarcinoma regulated by Msi2. Our functional experiments inhibiting these genes showed that loss of *Ptgds*, *Arl2bp*, *Rnf157*, and *Syt11* can block lung cancer sphere formation. Previous work has shown that Ptgds indirectly facilitates phosphorylated Sox9 nuclear translocation via activation of cAMP-dependent protein kinase A (PKA)^47^ and is important in the development of Sertoli cells. While *Ptgds* has been shown previously to be expressed in human lung tumors^58^ our work is the first to identify a functional role of Ptgds in lung adenocarcinoma. As a regulator of STAT3 signaling, Arl2bp directly binds to the STAT3 transcription factor, which maintains STAT3’s phosphorylation status and its localization in the nucleus^48^. STAT3 activity has been implicated in NSCLC^59^, thus modulating STAT3 activity through *Arl2bp* inhibition may provide an alternative therapeutic strategy. Lastly, both Rnf157 and Syt11 are predominantly expressed in the brain and have been implicated in neuronal cell survival and dendrite growth and neuronal vesicular trafficking, respectively^45,46^. Thus far, their roles have not been investigated in cancer. Our findings suggest the need for further study of these genes and raise the possibility that they could serve as future therapeutic targets.

While early treatment approaches centered principally on the use of cytotoxic agents, the treatment of lung cancer has evolved to incorporate the use of targeted therapies and immunotherapy, when appropriate, in combination with the standard cytotoxic regimen^60^. For patients with a lung cancer subtype harboring known targetable mutations, this has led to considerable improvements in the response to treatment. Yet troubling survival rates, driven largely by either late-stage detection or recurrence of the disease—even in cases for which a targeted therapy has been offered— illustrate the need for an increased understanding of the biology of lung cancer in order to inform future therapeutic approaches. The findings presented here identify a novel role for Msi2 in the growth and progression of lung adenocarcinoma and suggest that this role extends to human disease. Further, these findings identify novel effectors of lung adenocarcinoma regulated by Msi2 thereby lending new insight into the mechanism by which Msi2 exerts its influence on this disease. These new insights enhance our understanding of the biology of lung adenocarcinoma and may serve to open new avenues of focus for future therapeutic strategies.

## Methods

### Mice

Msi2^eGFP/+^ knock-in reporter mice were generated as previously described^6^ and are heterozygous for the Msi2 allele. The Msi2 mutant mouse, B6; CB-Msi2Gt(pU-21T)2Imeg (Msi2−/−) was established by gene trap mutagenesis as previously described^7^. The LSL-Kras G12D mouse, B6.129S4-Krastm4Tyj/J (stock number 008179), the p53flox/flox mouse, B6.129P2-Trp53tm1Brn/J (stock number 008462), and the *Rosa26*-*CreER^T2^* mouse, B6;*129-Gt*(*ROSA*) *26Sor^tm1(cre/Esr1)Tyj^* (stock number 008463) were purchased from The Jackson Laboratory. At 9 to 16 weeks of age 5X10^6^ PFU Adeno-Cre (purchased from the University of Iowa) was delivered by intratracheal instillation as previously described^25^. Where indicated, adult mice were administered tamoxifen (Sigma) in corn oil (20 mg/mL) daily by intraperitoneal injection (150 mg per gram of body weight) for 5 consecutive days. Lungs were then isolated 3 days following the last dose of tamoxifen. Mice were bred and maintained in the animal care facilities at the University of California San Diego. All animal experiments were performed according to protocols approved by the University of California San Diego Institutional Animal Care and Use Committee.

### Tissue Dissociation and FACS Analysis

For isolation of non-tumorigenic lung tissue, tissue dissociation was performed as described^61^. Briefly, mice were anesthetized, perfused with cold PBS using a peristaltic pump (Fisher Scientific), and 2mL dispase (Corning) was added to the lungs via intratracheal instillation. The lungs were then removed, placed on ice, and the lobes were minced. For isolation of lung tumors, mice were anesthetized and perfused with PBS as described, then lungs were removed and placed on ice. Visible tumors were dissected away from surrounding tissue, placed on ice in 2mL dispase, and minced. Minced non-tumorigenic lung tissue or grossly dissected lung tumors were then added to 2mg/mL collagenase/dispase in PBS (Millipore Sigma), placed in a rotisserie, and rotated for 45 minutes at 37°C. 25ng/mL DNAseI (Sigma-Aldrich) was added, and the tissue was then passed through a 100µm filter and a 40µm filter (Corning), and centrifuged at 1200rpm for 5 min at 4°C. Cells were then treated with 1mL red blood cell lysis buffer (Thermofisher) on ice for 90 seconds, washed, and resuspended in HBSS (Gibco, Life Technologies) containing 5% FBS and 2mM EDTA before staining. Analysis and cell sorting was performed using a FACSAriaIII machine (Becton Dickinson) and data were analyzed using FlowJo software (Tree Star). The following antibodies were used: rat anti-mouse EpCAM-APC (eBioscience), CD45-PE or CD45-PeCy7 (eBioscience), CD31-PE or CD31-PeCy7 (eBioscience), Sca1-PeCy7 (eBioscience). CD31 and CD45 antibodies were used to mark the Lineage^+^ populations of the lung. Specific antibody information is summarized in Supplementary Figure 2.

### Histology, Immunohistochemistry, and Immunofluorescence

For the isolation of tumor bearing lungs from KP^f/f^ mice, mice were anesthetized, perfused with PBS as described, and 4% PFA (Fisher Scientific) was added to the lungs via intratracheal instillation. Lungs were then removed, placed in 4% PFA overnight at 4°C, and stored in 70% ethanol prior to processing. The fixed lung tissue was then paraffin embedded at the UCSD Histology and Immunohistochemistry Core at The Sanford Consortium for Regenerative Medicine according to standard protocols, and 5 μm sections were obtained. Paraffin embedded tissues were then deparaffinized in xylene and rehydrated in a 100%, 95%, 70%, 50%, and 30% ethanol series. For immunohistochemistry, endogenous peroxidase was quenched in 3% hydrogen peroxide (Fisher Scientific) for 30 min at room temperature prior to antigen retrieval. Antigen retrieval was performed for 30 min in 95–100 °C 1× citrate buffer, pH 6.0 (eBioscience). Sections were blocked in TBS or PBS containing 0.1% Triton X100 (Sigma-Aldrich), 10% goat or donkey serum (Sigma Aldrich), and 5% bovine serum albumin. For single cell suspensions of lung epithelia or lung tumors, Lin^-^EpCAM^+^ cells were isolated via FACS in the manner described previously. Cells were resuspended in DMEM:F12 (Fisher Scientific) supplemented with 50% FBS and adhered to slides by centrifugation at 500rpm. 24 hours later, cells were fixed in 4% PFA (Fisher Scientific), washed in PBS, and blocked with PBS containing 0.1% Triton X-100 (Sigma-Aldrich), 10% Goat serum (Fisher Scientific), and 5% bovine serum albumin (Invitrogen).

All incubations with primary antibodies were carried out overnight at 4°C. For immunofluorescence staining, incubation with Alexafluor-conjugated secondary antibodies (Molecular Probes) was performed for 1 hour at room temperature. DAPI (Molecular Probes) was used to detect DNA and images were obtained with a Confocal Leica TCS SP5 II (Leica Microsystems). For immunohistochemical staining, incubation with biotinylated secondary antibodies (Vector Laboratories) was performed for 45 min at 20– 25°C. Vectastain© ABC HRP Kit (Vector Laboratories) was used according to the manufacturer’s protocol. Sections were counterstained with haematoxylin. The following primary antibodies were used: chicken anti-GFP (Abcam, ab13970), rabbit anti-Msi2 (Abcam, ab76148), rabbit anti-SPC (Santa Cruz, sc-13979), goat anti-CC10 (Santa Cruz, sc-9773). Specific antibody information is summarized in Supplementary Figure 3. Whole slide imaging was used to determine the number of tumors, tumor burden, and tumor grade in the tumor-bearing lungs of KP^f/f^ mice. H&E-stained slides were scanned using Aperio AT2 digital whole slide scanning (Leica Biosystems), and tumor analysis was performed using ImageScope (Aperio). Tumor grade was scored according to the criteria described by Jackson et al., 2005^53^.

### Mouse Lung Cancer Cell Lines

Mouse primary lung cancer cell lines were established from end-stage KP^f/f^ mice as follows: tumors were isolated and dissociated into single cell suspensions in the manner described above, then plated in 1X DMEM:F12 (Fisher Scientific) supplemented with 10%FBS, 1X pen/strep (Fisher Scientific), 1X N-2 supplement (Life Technologies), 35ug/mL bovine pituitary extract (Fisher Scientific), 20ng/mL recombinant mouse EGF (Biolegend), 20ng/mL recombinant mouse FGF (R&D Systems). Upon reaching 80% confluency the cells were collected and resuspended in HBSS (Gibco, Life Technologies) containing 5% FBS and 2mM EDTA, treated with FC block, and stained with EpCAM-APC (eBioscience), and CD31-PE (eBioscience). CD31-EpCAM+ cells were sorted and plated for at least one additional passage prior to use in functional assays. Functional assays were performed using cell lines between passage 2 and passage 6.

### Patient-Derived Xenografts

Patient pleural effusate samples were obtained from Moores Cancer Center at the University of California San Diego from Institutional Review Board-approved protocols with written informed consent in accordance with the Declaration of Helsinki. Effusates were centrifuged at 1200 rpm for 5 minutes at 4°C to pellet circulating tumor cells. Cells were then treated with 1mL red blood cell lysis buffer (Thermofisher) on ice for 90 seconds, washed in HBSS (Gibco, Life Technologies) containing 5% FBS and 2mM EDTA, and resuspended in 1X DMEM:F12 (Fisher Scientific) supplemented with 10%FBS, 1X pen/strep (Fisher Scientific), 1X N-2 supplement (Life Technologies), 35ug/mL bovine pituitary extract (Fisher Scientific), 20ng/mL recombinant human EGF (Life Technologies), 20ng/mL recombinant human FGF-basic (Fisher Scientific). For each patient sample, a 1:1 ratio of resuspended cells in growth factor reduced Matrigel (Corning) at a final volume of 100 µL was transplanted into the flank NSG recipient mice to allow for the development of the patient-derived xenograft. Tumor growth was monitored via caliper measurement, and mice were euthanized when tumors reached 2 cm in diameter. Tumors were then dissociated and used for functional analysis or for subsequent passage. Functional analysis was performed using patient-derived xenografts between passage 1 and passage 2.

### Lung Tumorsphere Assays

For tumorsphere assays of mouse lung cancer cell lines involving genetic inhibition via shRNA, low passage KP^f/f^ cell lines were infected with shRNA lentivirus. 72-hours after transduction, the cells were sorted for positively infected cells. For tumorsphere assays involving Msi2^eGFP/+^; Kras^G12D/+^; p53^fl/fl^; RosaCre^ER/+^ mice, tamoxifen treatment and cell sorting for CD45-CD31-EpCAM+GFP+ (Msi2^+^) and CD45-CD31-EpCAM+GFP-(Msi2-) cell populations were performed as described. After sorting, cells were resuspended in full media [1X DMEM:F12 (Fisher Scientific) supplemented with 10%FBS, 1X pen/strep (Fisher Scientific), 1X N-2 supplement (Life Technologies), 35ug/mL bovine pituitary extract (Fisher Scientific), 20ng/mL recombinant mouse EGF (Biolegend), 20ng/mL recombinant mouse FGF (R&D Systems)]. Resuspended cells were mixed with an equal volume of growth factor reduced Matrigel (Corning) for a total volume of 100µL, plated in 96 well flat-bottom plates (Fisher Scientific) at a density of 500 cells/well. Matrigel was allowed to solidify at 37°C for 20 min, and 150 µL of full media was added to the well. Tumorsphere cultures were incubated at 37°C for 10-14 days with the media refreshed once a week. To passage cells, tumorspheres were dissociated from surrounding Matrigel by removing the media, adding 200 µL of 2mg/mL collagenase/dispase (Millipore Sigma) to the wells, and incubating for 1 hour at 37°C. Cells were collected and placed on ice, the wells were washed 3X with full media, and each media collection was pooled on ice with the original cell collection. Cells were pelleted via centrifugation at 500 rcf for 5 minutes at 4°C, media was aspirated, 200 µL of cold Accumax (Innovative Cell Technologies) was added to the cell pellet, and cells were incubated at 37°C for 10 minutes. Cells were manually pipetted 100X to dissociate tumorspheres and examined via hemocytometer to determine if a single cell suspension had been obtained. The incubation and mechanical dissociation were repeated 2-3X until a single cell suspension was obtained. Cells were resuspended in full media, and 500 cells were plated at a 1:1 ratio with growth factor reduced Matrigel (Corning) at a final volume of 100 µL. Matrigel was allowed to solidify at 37°C for 20 min, and 150 µL of full media was added to the well, and cells were incubated in the manner described.

### Flank Transplant Assays

For flank transplant assays of mouse lung cancer cell lines involving genetic inhibition via shRNA, low passage KP^f/f^ cell lines were transduced with shRNA lentivirus, sorted, and 20,000 cells were resuspended in a 1:1 mix of full media and Matrigel (Corning) at a final volume of 100 µL in the manner described. Cells were injected subcutaneously into the flank of non-tumor bearing (i.e. not treated with Adeno-Cre), immunocompetent KP^f/f^ mice. Subcutaneous tumors were measured with calipers once weekly for 6-15 weeks and did not exceed 2cm in diameter. At endpoint, flank tumors were removed, and tumors were dissociated as described. Tumor cells were stained with CD45-PeCy7 and CD31-PeCy7, and the absolute number of tumor cells was calculated by multiplying the percentage of CD45-CD31- transduced cells by the total number of CD45-CD31- live cells. For flank transplant assays of patient-derived xenografts, freshly dissociated xenografts were plated in full media [1X DMEM:F12 (Fisher Scientific) supplemented with 10%FBS, 1X pen/strep (Fisher Scientific), 1X N-2 supplement (Life Technologies), 35ug/mL bovine pituitary extract (Fisher Scientific), 20ng/mL recombinant human EGF (Life Technologies), 20ng/mL recombinant human FGF-basic (Fisher Scientific)]. 500,000 cells/well were plated in a 6-well plate (Fisher Scientific) coated with Matrigel (Corning) for subsequent transplant, and 83,000 cells/well were plated in a 24-well plate (Fisher Scientific) coated with Matrigel (Corning) for subsequent determination of transduction efficiency via FACS. Cells were transduced with proportionate amounts of shControl or shMsi2 lentivirus. 24 hours after transduction, media from the cells intended for transplant was collected and placed on ice. 1mL of 2mg/mL collagenase/dispase (Millipore Sigma) was then added to the well and incubated for 45 minutes at 37°C to dissociate the remaining cells from Matrigel. The wells were washed 3X with full media and each wash was collected and pooled on ice with the media previously collected. Cells were pelleted by centrifugation at 1200 rpm for 5 minutes at 4°C, an equivalent number of shControl and shMsi2 transduced cells were resuspended in full media, mixed at a 1:1 ratio with growth factor reduced Matrigel (Corning) at a final volume of 100 µL, and transplanted subcutaneously into the flanks of NSG recipient mice. Tumor growth was monitored 1-2X a week via caliper measurement for 11-12 weeks and did not exceed 2cm in diameter. 48 hours after transduction, cells from the 24-well plate were collected in the manner described using a proportionate amount of reagents, and the frequency of transduced live cells was determined via FACS. For flank transplant assays of cells from Msi2^eGFP/+^; Kras^G12D/+^; p53^fl/fl^; RosaCre^ER/+^ tumorspheres, quaternary passage tumorspheres were dissociated from surrounding Matrigel in the manner described. Cells were resuspended in full media, mixed at a 1:1 ratio with growth factor reduced Matrigel (Corning) at a final volume of 100 µL, and transplanted subcutaneously into the flanks of NSG recipient mice. Tumor growth was monitored 1-2X a week via caliper measurement for 13 weeks and did not exceed 2cm in diameter. At endpoint, both the flank tumor and lungs were collected for fixation and H&E analysis in the manner described. Lung H&E sections were analyzed for the presence of metastatic tumors.

### shRNA Lentivirus Production

For knockdown of Msi2, shRNA constructs and scrambled control were designed as previously described^6^. For knockdown of Ptgds, Arl2bp, Rnf157, and Syt11, shRNA constructs and scrambled control were designed as previously described^62^ using the following target sequences: For Ptgds, 5’-ACCTCTACCTTCCTCAGGAAA – 3’; for Arl2bp, 5’-GCTGCTCACATTCACGGATTT-3’; for Rnf157 5’-CAGAGGGAAATGATATCATAG – 3’; for Syt11, 5’-ATCAGGCTTCTCTGGGTTATT-3’.

### qRT-PCR Analysis

RNA isolation was performed using RNeasy Micro and Mini kits (Qiagen) and cDNA conversion was performed using Superscript III (Invitrogen). Quantitative real-time PCR was performed using an iCycler (BioRad) in a mix containing iQ SYBR Green Supermix (Biorad), cDNA, and gene-specific primers. Primer sequences for target genes are shown in Supplementary Figure 3. All real-time data was normalized to B2M.

### RNA sequencing and Bioinformatics Analysis

For RNA-seq of lung epithelial cells from Msi2^eGFP/+^; Kras^G12D/+^; p53^fl/fl^; RosaCre^ER/+^ mice, 60,000 cells were isolated for the Msi2^+^ (Lin-EpCAM+GFP+) and Msi2^-^(Lin-EpCAM+GFP) populations via FACS in the manner described. For RNA-seq of Msi2 knockdown in KP^f/f^ cells, KP^f/f^ cells were transduced with either shControl or shMsi2 lentivirus in triplicate for each group, and ≥125,000 transduced cells were isolated via sorting. RNA was isolated using a RNeasy Micro Kit (Qiagen). The quality of total RNA was assessed using an Agilent Tapestation and all samples had a RIN ≥8. M2KD RNA libraries were generated from 2 µg of RNA, and Msi2^eGFP/+^; Kras^G12D/+^; p53^fl/fl^; RosaCre^ER/+^ libraries were generated from 90ng of RNA using Illumina’s TruSeq Stranded mRNA Sample Prep Kit following the manufacturer’s instructions. RNA libraries were multiplexed and sequenced with 50 basepair (bp) single end reads (SR50) to a depth of approximately 30 million reads per sample on an Illumina HiSeq4000.

RNA-seq analysis and cell state analysis were performed as previously described^62^. For validation of differential expression analysis, edgeR and sleuth were used. Briefly, RNA-seq fastq files were processed within the systemPipeR pipeline^63^. Alignment of the reads was performed using HISAT2 with the mm10 mouse assembly using default settings^64^. Read counting was performed with the summarize Overlaps R packages with Union mode and the edgeR package was used for differential expression analysis using a fold change >1.9 and an FDR <0.05^65,66^. As a secondary validation, transcript quantifications were performed using kallisto and sleuth was used for gene-level quantifications and differential expression analysis using default setting^67,68^.

### Statistical Analysis

Statistical analysis was performed using GraphPad Prism software version 7.0 (GraphPad Software Inc.). Data are mean ± SEM. Two-tailed unpaired Student’s t-tests and Log-Rank tests were used to determine statistical significance. The Grubb’s test was used to identify and remove statistical outliers where indicated.

## Data Availability

The datasets generated and analyzed during the current study are available from the corresponding author on reasonable request.

## Acknowledgements

We are grateful to Reuben J. Shaw for scientific advice; Laurie Gerkin and Amanda Hutchins for technical training; Lillian J. Eichner for technical advice; and Marcie Kritzik for assistance with manuscript preparation.

---

**Supplementary Figure 1:**
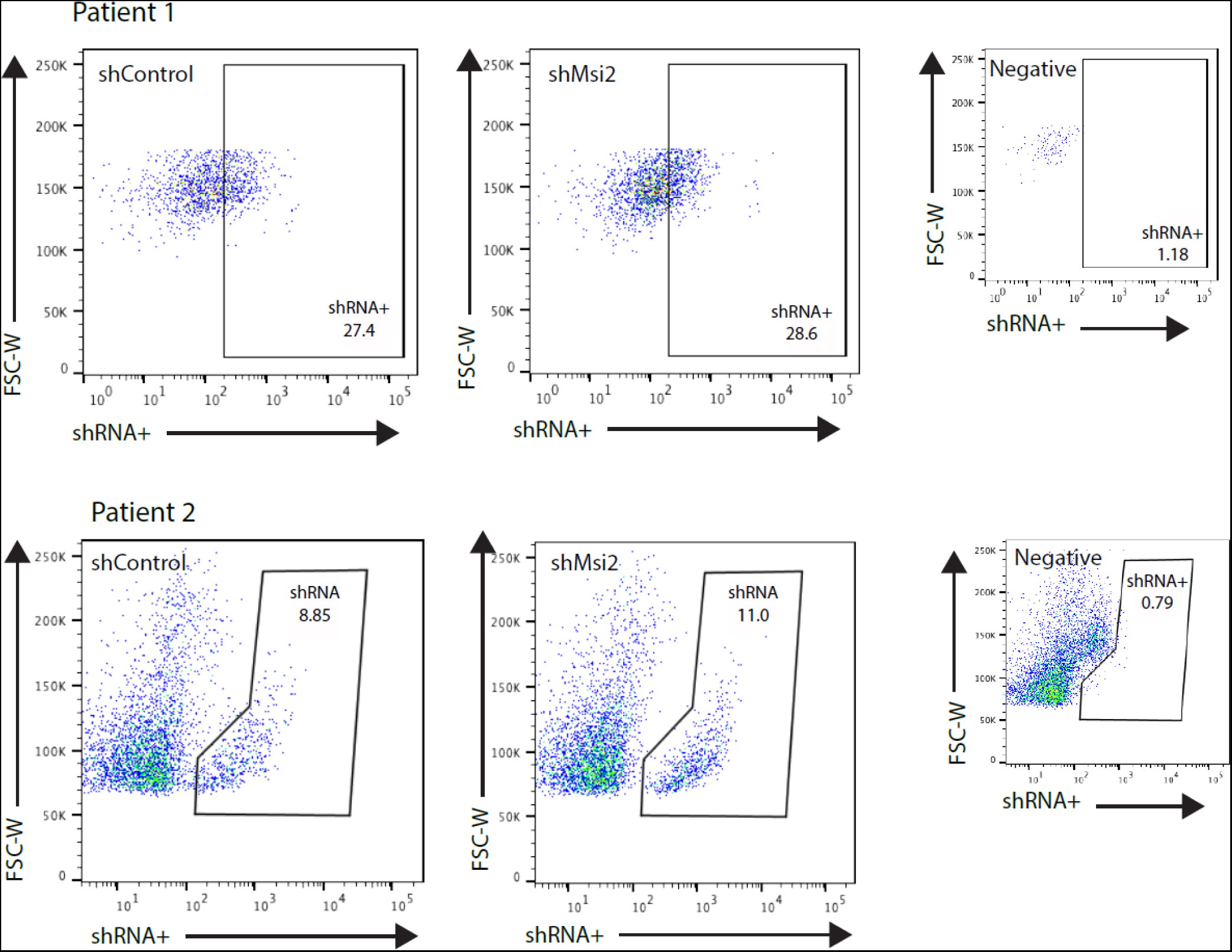
FACS plots for PDXs transduced with shRNA. PDX cells transduced with either shControl or shMsi2 lentivirus have a similar frequency of infection. CD45-CD31-PDX cells from Patient 1 show a 27.4% frequency of infection when transduced with shControl, and a 28.6% frequency of infection when transduced with shMsi2 (top). CD45-CD31-PDX cells from Patient 2 show an 8.85% frequency of infection when transduced with shControl, and an 11% frequency of infection when transduced with shMsi2 (bottom). PDX cells that were untransduced and unstained for FACS antibodies were used as negative gating controls (shown at right for each patient sample).

**Supplementary Figure 2:**
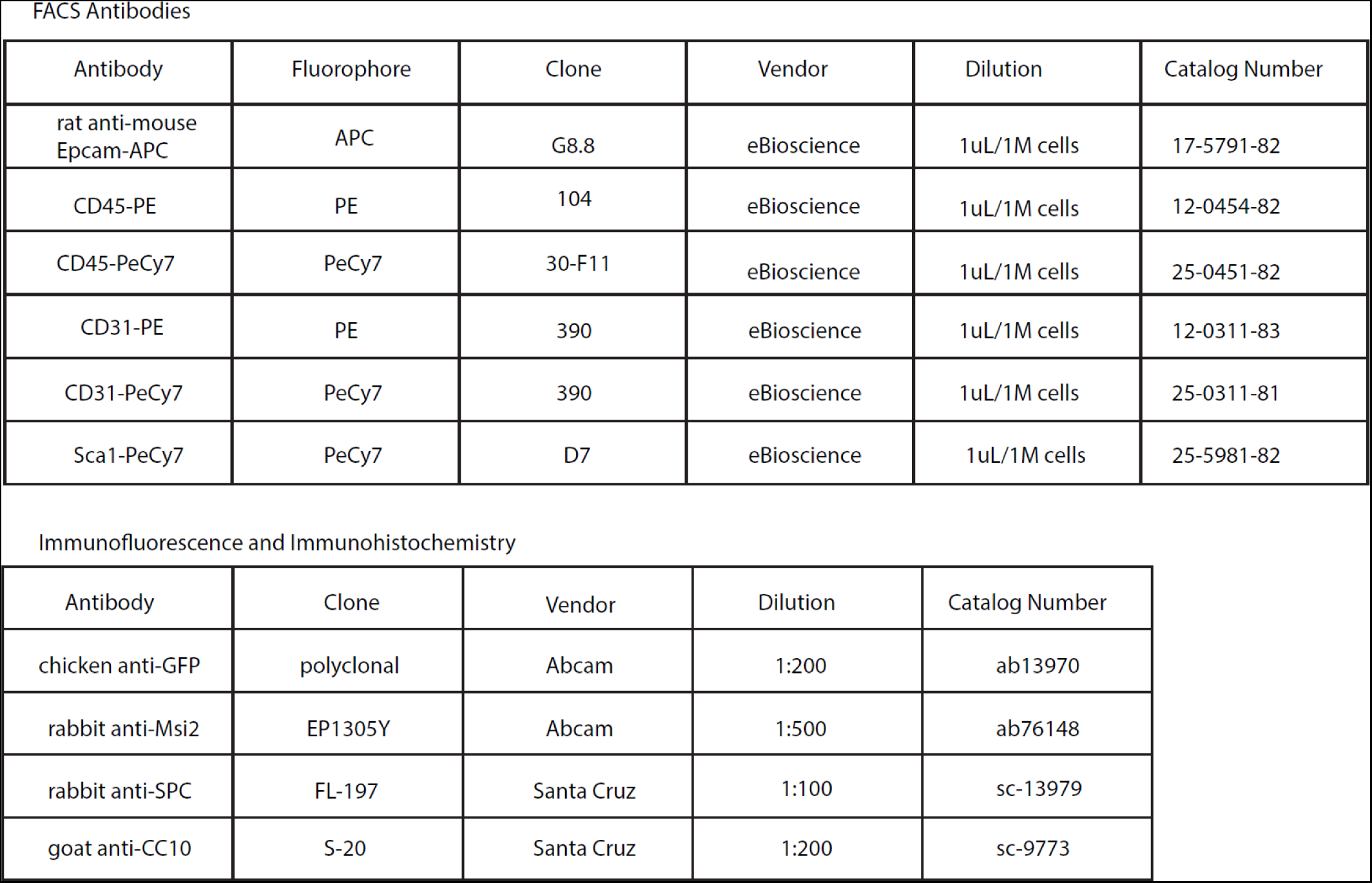
Antibodies used Tables of antibodies used in this study for FACS and immunofluorescence and immunohistochemistry.

**Supplementary Figure 3:**
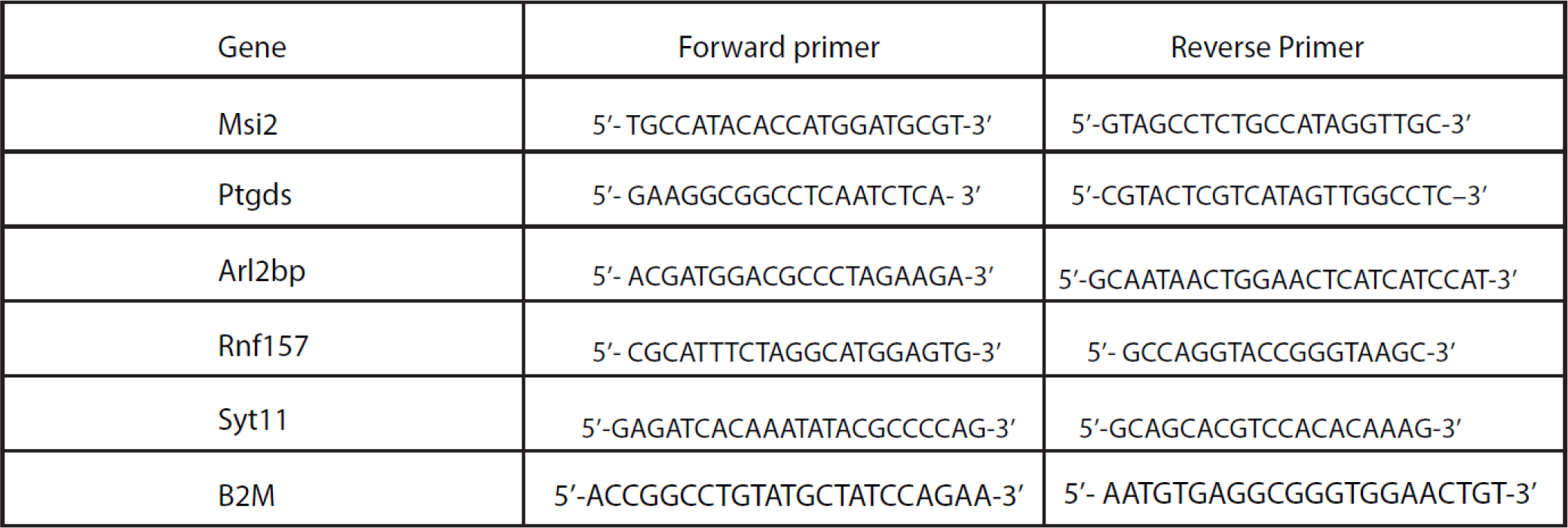
qRT-PCR primers for genes of interest Table of qRT-PCR primers for all genes of interest as well as the housekeeping gene used in this study.

